# Triceps surae torque-length relationships relevant for walking activity levels with and without an ankle exoskeleton

**DOI:** 10.1101/778175

**Authors:** Anthony L. Hessel, Brent J. Raiteri, Michael J. Marsh, Daniel Hahn

## Abstract

**Abstract:** Ankle exoskeletons have been developed to assist walking by offloading the plantar flexors work requirements, which reduces muscle activity level. However, reduced muscle activity alters plantar flexor muscle-tendon unit dynamics in a way that is poorly understood. We therefore evaluated torque-fascicle length properties of the soleus and lateral gastrocnemius during voluntary contractions at simulated activity levels typical during late stance with and without an ankle exoskeleton. Soleus activity levels (100, 30, and 22% maximal voluntary activity) were produced by participants via visual electromyography feedback at ankle angles ranging from −10° plantar flexion to 35° dorsiflexion. Using dynamometry and ultrasound imaging, torque-fascicle length data of the soleus and lateral gastrocnemius were produced. The results indicate that muscle activity reductions observed with an exoskeleton shift the torque-angle and torque-fascicle length curves to more dorsiflexed ankle angles and longer fascicle lengths where no descending limb is physiologically possible. This shift is in line with previous simulations that predicted a similar increase in the operating fascicle range when wearing an exoskeleton. These data suggest that a small reduction in muscle activity causes changes to torque-fascicle length properties, which has implications for the design and testing of future ankle exoskeletons for assisted walking.

**Significance Statement:** Assistive lower-limb exoskeletons reduce the metabolic cost of walking by reducing the positive work requirements of the plantar flexor muscles. However, if the exoskeleton reduces plantar flexor muscle activity too much, then the metabolic benefit is lost. The biological reasons for this are unclear and hinder further exoskeleton development. This research study is the first to directly evaluate if a reduction in plantar flexor muscle activity similar to that caused by wearing an exoskeleton affects muscle function. We found that reduced muscle activity changes the torque-length properties of two plantar flexors, which could explain why reducing muscle activity too much can increase metabolic cost.

## Introduction

Human walking is the most utilized energy-consuming activity for healthy humans and is closely associated with a person’s quality of life and the prevalence of secondary disease states (Tudor-Locke et al., 2011). Walking is remarkably adaptable to changes in terrain, such as slopes (Alexander et al., 2017; Lay et al., 2006; Pickle et al., 2016), stairs (Lewis et al., 2015), surface compliances (MacLellan and Patla, 2006), and unexpected perturbations (Vlutters et al., 2018) because of robust neural control strategies and intrinsic muscle properties (Clark, 2015).

Many research groups fabricate anthropometric wearable devices for walking, called exoskeletons, which fit closely to the lower-limb of the body and function in concert with the user’s limbs to augment and assist movement. Of interest for this study is the rapid development of passive, non-motorized, lower-limb exoskeletons. The industry’s measuring stick to validate the benefit of these assistive devices to the user is a reduced metabolic cost of transport for a healthy user at a preferred walking speed on level ground (Herr, 2009). By this standard, the design by Collins et al. Collins et al. (2015) is currently the most successful passive lower-limb exoskeleton. The exoskeleton design employs a bio-inspired spring-clutch system that places a linear tension spring in parallel with the plantar flexor muscles in such a way that the spring stores and releases elastic energy along with the plantar flexor muscle-tendon units (MTUs) during the stance phase of walking (Collins et al., 2015). This ankle exoskeleton reduces the metabolic cost of transport during steady, single-speed, level ground walking by ~7.2% compared with unassisted walking (walking without an exoskeleton), presumably by off-loading some of the work requirements of the plantar flexors during the stance phase of walking, which is supported by a measured reduction in soleus and gastrocnemii muscle activities when using the exoskeleton (Collins et al., 2015). Interestingly, future improvements have been minimal because off-loading the muscles too much (i.e. by increasing the exoskeleton spring stiffness) leads to *increased* metabolic costs caused by complex human-machine interactions (Jackson et al., 2017; Steele et al., 2017) that are poorly understood.

Several research groups have begun to define these complex human-machine interactions when wearing a lower-limb exoskeleton in order to outline current ankle exoskeleton shortcomings and to provide a foundation for the optimization of future devices. One research focus is on the interaction between MTU dynamics and the reduction in muscle activity when using a lower-limb exoskeleton. The plantar flexor MTUs properties, such as muscle activity patterns and tendon stiffness, are highly tuned to optimize muscle efficiency during walking (Lichtwark and Wilson, 2008), so small changes to activity level could have large consequences. For example, two modeling studies (Jackson et al., 2017; Sawicki and Khan, 2016) suggest that increasing the amount of exoskeleton torque (i.e. by increasing the spring stiffness in the Collins et al. (Collins et al., 2015) design changes the plantar flexors MTU mechanics and reduces the plantar flexors positive power output. Thus, these overall losses in MTU performance negate the positive benefits of reduced muscle activity level at a certain threshold of muscle unloading.

Another possible consequence of reduced muscle activity levels on muscle and locomotor performance is related to where muscles operate on their force-length curves. MTU compliance and activation-dependent shifts in optimal fascicle length (Holt and Williams, 2018; Ichinose et al., 1997; MacIntosh, 2017) can increase the fascicle length that maximal force is produced with decreasing activity levels. Because plantar flexor muscle activity is reduced with exoskeleton assistance, it is likely that optimal fascicle length is also affected, perhaps adding to the performance deficits associated with exoskeleton assistance described above. Unfortunately, little is known about activation-dependent changes of the triceps surae fascicle length operating ranges for muscle activities relevant for walking with and without an exoskeleton.

The aim of this study was to evaluate the activation-dependence of triceps surae optimal fascicle length to better understand whether exoskeleton use might change the torque-producing capabilities of the major plantar flexors compared with normal human walking. We studied the soleus (SOL) and lateral gastrocnemius (LG) muscles during fixed-end voluntary contractions at maximal effort and at submaximal voluntary activity levels typical for soleus for walking with and without an exoskeleton (15). We focused on identifying the net (plantar flexion) ankle torque–angle and torque–fascicle length relationships. We hypothesized that decreased activity levels would shift the ankle joint torque-ankle angle and ankle joint torque–fascicle length relationships towards more dorsiflexed ankle angles and longer fascicle lengths.

## Methods

### Participants and Ethics

Healthy male and female participants (n = 8/4 male/female, age = 25.5 ± 2.4 years, weight = 77.9 ± 11.2 kg, height 177.4 ± 5.6 m) were recruited from Ruhr University Bochum. Participants were free of neuromuscular disorders or injuries to their lower limbs and had no documented gait abnormalities.

### Experimental Setup

#### Dynamometer

Net ankle joint torque and ankle joint angle were measured from the right leg of participants using an isokinetic dynamometer (Fig. 1; IsoMed2000, D&R Ferstl GmbH, GER). Participants laid prone on the bench of the dynamometer with their knee slightly bent (~5°) and their foot tightly strapped to a foot plate to avoid heel lift during contractions. Further setup details available in Suppl. Info. 1.

**Figure 1:**
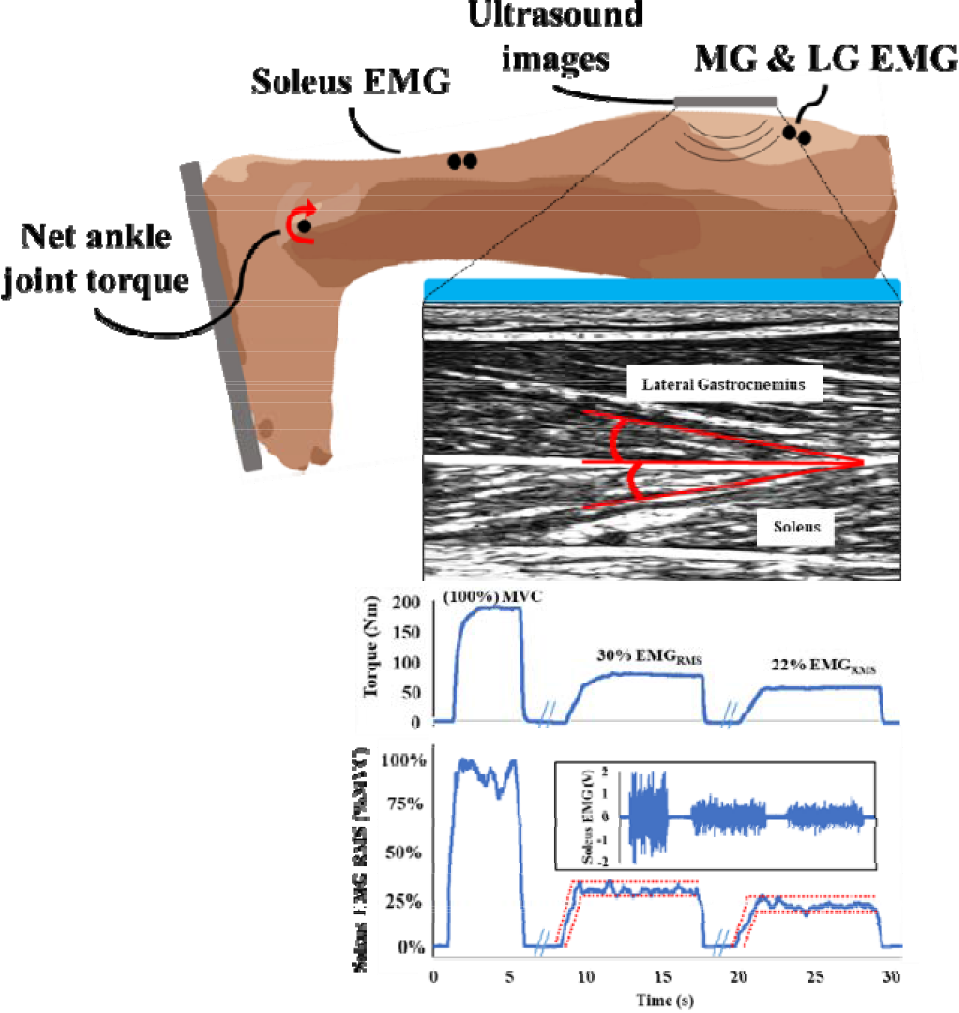
Representative example of the *in vivo* dynamometer study. 30% and 22% SOL activity levels were EMG controlled via bio-feedback. Top: EMG electrodes were placed on the SOL, MG and LG. An ultrasound probe was placed on the lateral gastrocnemius in such a way that both LG and SOL fascicles were imaged. Middle: Ultrasound images were used to measure fascicle length. Bottom: SOL EMG activity was voluntarily controlled by the subjects as they watched real-time feedback of their SOL EMG RMS amplitude averaged over 250 ms. To guide the subject, target activity zones were drawn onto the screen (red dotted lines). Torque data during EMG-controlled experiments was consistent at the same EMG level.

#### Surface electromyography

Electromyography (EMG) was used to measure the muscle activity of the right leg’s soleus (SOL) and lateral and medial gastrocnemii (LG and MG) (AnEMG12, OT Bioelettronica, Torino, Italy). For easy visualization, raw EMG signals had the DC offset removed and were smoothed with a 0.25◻second root-mean-square (RMS) amplitude calculation. Further details available in Suppl. Info. 2.

#### Ultrasound measurements

An 128-element flat, linear ultrasound transducer (60 Hz sampling rate, 60 × 50 mm (width × depth) range view, LS128 CEXT-1Z, Telemed, Lithuania) was used to image SOL and LG fascicles simultaneously throughout all experiments, as detailed elsewhere (Farris et al., 2016). Each muscle’s fascicle lengths were measured in each image from a representative fascicle using already described tracking software and procedures (Farris and Lichtwark, 2016; Gillett et al., 2013).

#### Activity level selection and control

The SOL muscle activity levels were designed to replicate those observed during the mid-stance phase of level ground walking at a self-selected speed while walking with or without an assistive lower-limb exoskeleton (Collins et al., 2015). Selecting only two activity levels is difficult because EMG levels continue to change over the course of the stance phase, with or without an exoskeleton (Jackson and Collins, 2015). The average SOL EMG activity level over a whole stride is ~32% of the peak activity over a whole stride without the exoskeleton (see Extended Data Fig. 4; Collins et al., 2015). We normalized this value to maximal voluntary contraction (MVC) using data from Nishijimi e al. (Nishijima et al., 2010), where peak SOL EMG over a whole stride was ~95% MVC, which resulted in ~30.4% MVC. When using an exoskeleton, Collins et al. (Collins et al., 2015) reported an average reduction of ~22% during early- to mid-stance (0% – 40% of the whole stride). However, based on data from (Extended Data Fig. 4; Collins et al., 2015), this average reduction from heel-strike to mid-stance still underestimates the more typical reduction during mid-stance (ranging from ~25 – 29% reduction) because early-stance produces little to no EMG reduction. Thus, we selected a more typical reduction level observed during mid-stance. Taking the above considerations into account, controlled activity levels were set at 30% and 22% MVC of the peak-to-peak SOL EMG RMS amplitude during MVC (26.7% EMG reduction). EMG level validation is available (Suppl. Info. 1).

Participants received real-time biofeedback of their SOL activity level via a computer monitor so that target activity levels (22% and 30% MVC) could be achieved voluntarily for a 10 second test period, similar to experiments on the tibialis anterior (Raiteri and Hahn, 2019). The procedure required participants to match their real◻time SOL EMG (moving 0.25◻ second RMS amplitude) to within ± 5% of a predefined EMG RMS amplitude (Fig. 1). After a familiarization session, subjects became consistent at matching their EMG activity to the desired level.

### Experimental Protocol

#### Mechanical tests

The subject performed preconditioning contractions before experimental tests by completing at least 10, 3-5 s contractions that gradually increased in strength from ~50% to 100% perceived effort. The subject warmed up more if desired. At an ankle angle of 0° (sole of the foot perpendicular to the shank), subjects plantar flexed with maximal effort for 6 seconds to produce a MVC. Subjects were given verbal motivation and performed at least two trials with no more than 5% difference in peak torque. 3-minutes of rest was provided between contractions. The soleus RMS EMG level produced during the MVC that produced the largest plantar flexion torque was used to calculate the two target activity levels (30% MVC, 22% MVC). In total, subjects completed experiments at three activity levels: 22%, 30%, and 100%, and data was also collected when subjects were instructed to relax, which we will define as 0% activity. We quantified the activity-dependence of the torque-ankle angle, fascicle length-ankle angle, and torque-fascicle length relationships at 22%, 30%, and 100% activity levels. We measured the net ankle joint torque produced at each activity level at ankle angles of −10° (plantar flexion), 0°, 5° (dorsiflexion), 10°, 15°, 20°, 25°, 30°, and 35°. Dorsiflexion > 25° ankle angle was not a dorsiflexion range all subjects could reach without discomfort. Therefore, some of the extreme dorsiflexion positions were captured only for a subset of participants (n = 11 at 30°, n = 10 at 35°). Passive (0% activity level) torque data was also captured by moving the ankle between −10° and 35° at 1°s^−1^ for five full cycles and the last 2 rotations were averaged. The first three rotations were completed to precondition the subject to keep their muscle activity off during rotations.

Contractions were produced from the lowest to highest activity levels and the order of angles within each activity level was randomized. This procedure was selected to minimize the effects of fatigue from the MVCs. Each contraction lasted six and ten seconds for MVC and submaximal activity levels, respectively. Subjects received 3-minutes rest between MVC contractions, at least 30 seconds rest between submaximal activity level contractions, and 5-minutes rest between the activity conditions. More time was given upon request from the participant. To monitor fatigue, periodic MVCs were measured and testing concluded if torque was < 90% initial MVC. All torque data was normalized to %MVC at maximum torque, among the ankle angles tested. Each condition was repeated twice and averaged, unless the two torque values were >5% different (with successful EMG control), which resulted in a third measurement and an average being taken. During all trials, ultrasound videos of the LG and SOL fascicles were captured, and fascicle lengths were measured from those videos during the last 3 seconds of the steady-state contractions.

#### Data analysis

Active torque – ankle angle, active fascicle length – ankle angle, and active torque – fascicle length relationships were created from the data. Each data point during contraction was the average of the last 3 s of the contraction window. During 22% and 30% activity levels, subjects produced a steady-state activity level after ~5 seconds. 100% MVC reached a steady state within ~1 s. The separation of active torque from net torque was done by considering changes to passive torque caused by fascicle shortening during activation (Suppl. Info. 3).

### Statistical analysis

Active torque-ankle angle and active torque-active fascicle length relationship data were calculated for each participant, normalized to maximal torque, plotted with all participants, and a second-order polynomial equation was then fitted for each activity level. We used these equations to estimate optimal ankle angle and fascicle length at peak plantar flexion torque for the 100% activity level.

Muscle activity (MG/LG/SOL) and torque levels was normalized to their respective values during 100% MVC at 0°. To compare torque and EMG at different activity levels and angles, for each muscle, we designed a full-factorial, 2-way ANOVA with fixed effects for activity level (100%, 30%, 22%) and ankle angle (−10°, 0°, 5°, 10°, 15°, 20°, 25°, 30°, and 35°), and a random effect for the individual. Response parameters included torque and MG/LG/SOL activity levels. To compare the interaction between ankle angle, activity, and fascicle length on torque, we added a fascicle length covariate to our ANOVA model above, with a response parameter of torque. Alpha values were set at 0.05 and assumptions of normality and homogeneity of variance were evaluated using the Shapiro-Wilk test of normality and Levene’s test for equality of variances. To limit data skewing, outliers (data points > 2 standard deviations from the mean) were removed. A best Box-Cox transformation was used for all data sets to meet assumptions. When model effects were significant, a *post-hoc* Tukey Honestly Significant Difference (HSD) all-pairwise comparison analysis was used to test for differences among group means. Data are presented as mean ± standard error mean (SEM). All fitted lines on graphs show a 95% confidence interval of the fit. Statistical analysis was conducted using JMP (JMP Pro 12.2, SAS Institute Inc., Cary, NC, USA).

## Results

### Fascicle Length

SOL (Fig. 2A; Table 1) and LG (Fig. 2B; Table 1) fascicle lengths showed similar trends, with increasing lengths as the ankle moved to more dorsiflexed positions and decreasing lengths with increasing activity levels. During fixed-end contractions, SOL fascicles shortened on average 20.6%, 25.7% and 34.0% of passive length for the 22%, 30%, and 100% activity levels, respectively. LG fascicles shortened on average 21.6%, 23.5%, and 28.6% of passive length for the 22%, 30%, and 100% activity levels, respectively. For both muscles, 100% and 0% activity levels produced the shortest and longest fascicles, respectively (Tukey HSD, P < 0.05). However, only SOL fascicle lengths significantly decreased from 22% to 30% activity (Tukey HSD, P < 0.05). However, on visual inspection of LG data from each participant, a consistent decrease in fascicle length was present from the 30% to 22% activity levels at all angles.

**Figure 2:**
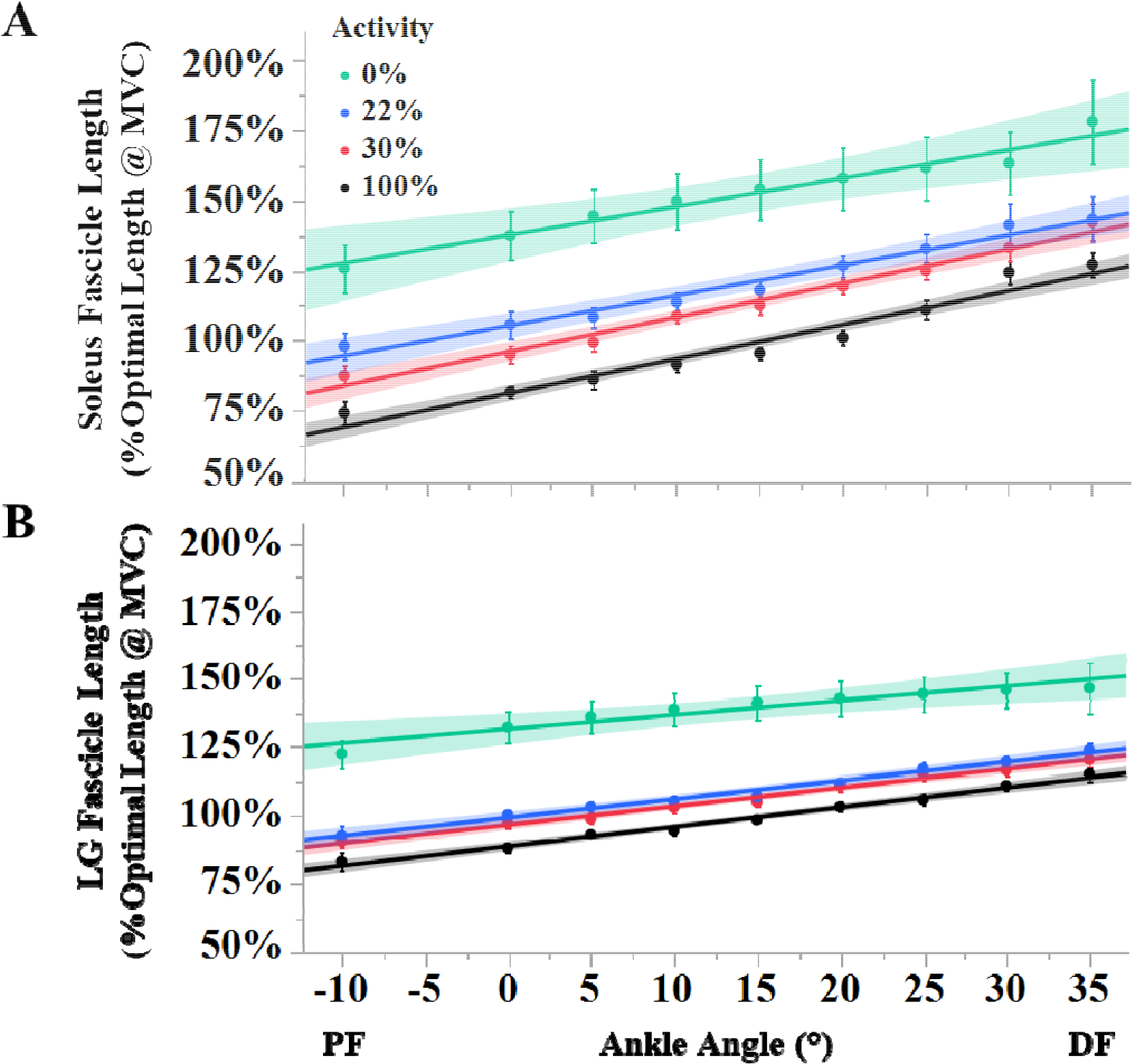
Fascicle lengths for SOL (A) and LG (B) at 0%, 22%, 30%, and 100% activity levels. Both muscles produced consistently shorter fascicle lengths at 22% activity compared to 30% activity, but only SOL was significantly different. A linear fit is drawn at each activity level and shaded regions indicate 95% confidence intervals of the fit. Data is normalized to the fascicle length at the ankle angle that produced the maximum torque during MVCs (100% activity). Error bars of data points are s.e.m.

**Table 1.**
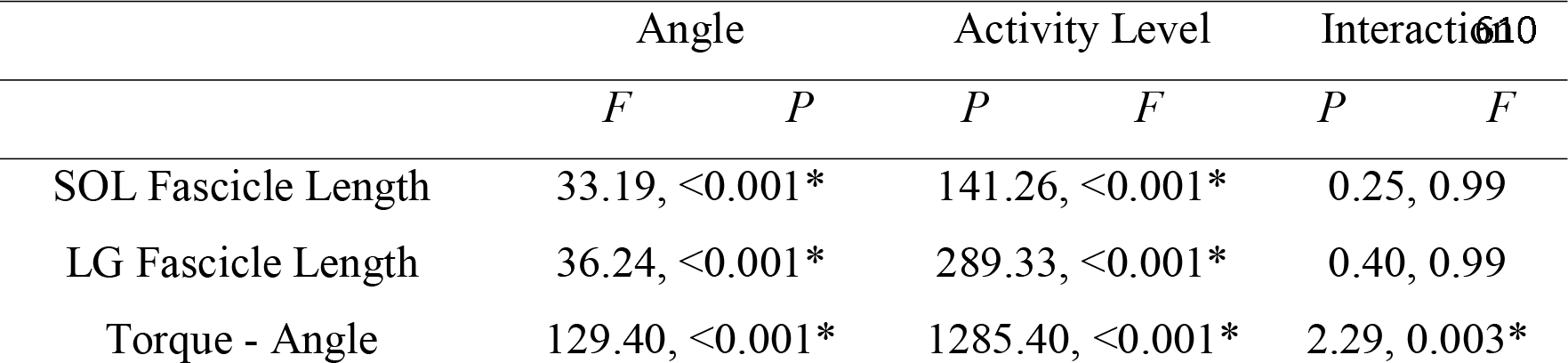
Results of two-way ANOVA test on plantar flexor MTU dynamics at different EMG activity levels. *F* = F statistic, *P* = P-value. * indicates significance (*P* < 0.05). Note: individual random response added to the model, and significant for all tests (Wald test, P < 0.05). SOL = soleus, LG/MG = lateral and medial gastrocnemius.

|  | Angle |  | Activity Level |  | Interaction |  |
| --- | --- | --- | --- | --- | --- | --- |
| | $F$ | $P$ | $P$ | $F$ | $P$ | $F$ |
| SOL Fascicle Length | 33.19, <0.001* |  | 141.26, <0.001* |  | 0.25, 0.99 |  |
| LG Fascicle Length | 36.24, <0.001* |  | 289.33, <0.001* |  | 0.40, 0.99 |  |
| Torque - Angle | 129.40, <0.001* |  | 1285.40, <0.001* |  | 2.29, 0.003* |  |

### Torque-ankle Angle Relationship

Net ankle joint torque decreased with decreasing muscle activity levels at all ankle angles (Tukey HSD, P < 0.05; Fig. 3; Table 2). At maximum torque, 22% and 30% activity levels produced ~48% and ~56% of the 100% MVC torque, respectively. Furthermore, both 22% and 30% activity levels produced a significantly larger amount of the 100% torque (at a given angle) as the ankle became more plantar flexed (Fig. 3; Table 2). The 100% activity level produced a typical torque-angle relationship, with a long ascending limb and peak active torque being produced at an estimated optimal angle of ~25° (Fig 3, vertical line). The statistical analysis indicates that values between 20° and 30° ankle angle were not different (note: individual variation taken into account, see methods), indicating a 10° wide plateau region during MVC and a small descending limb. For the 22% and 30% activity levels, there was no statistical difference between 25° and 35°, indicating a ~10° plateau region of the torque-angle relationship and no descending limb. Although it is clear that decreasing activity levels shifted the optimal ankle angle to more dorsiflexed positions via qualitative evaluation of the data, because no descending limb of the torque-ankle angle relationship was found for the 22% and 30% activity levels, the exact locations of the optimal ankle angles cannot be determined for the submaximal conditions.

**Figure 3:**
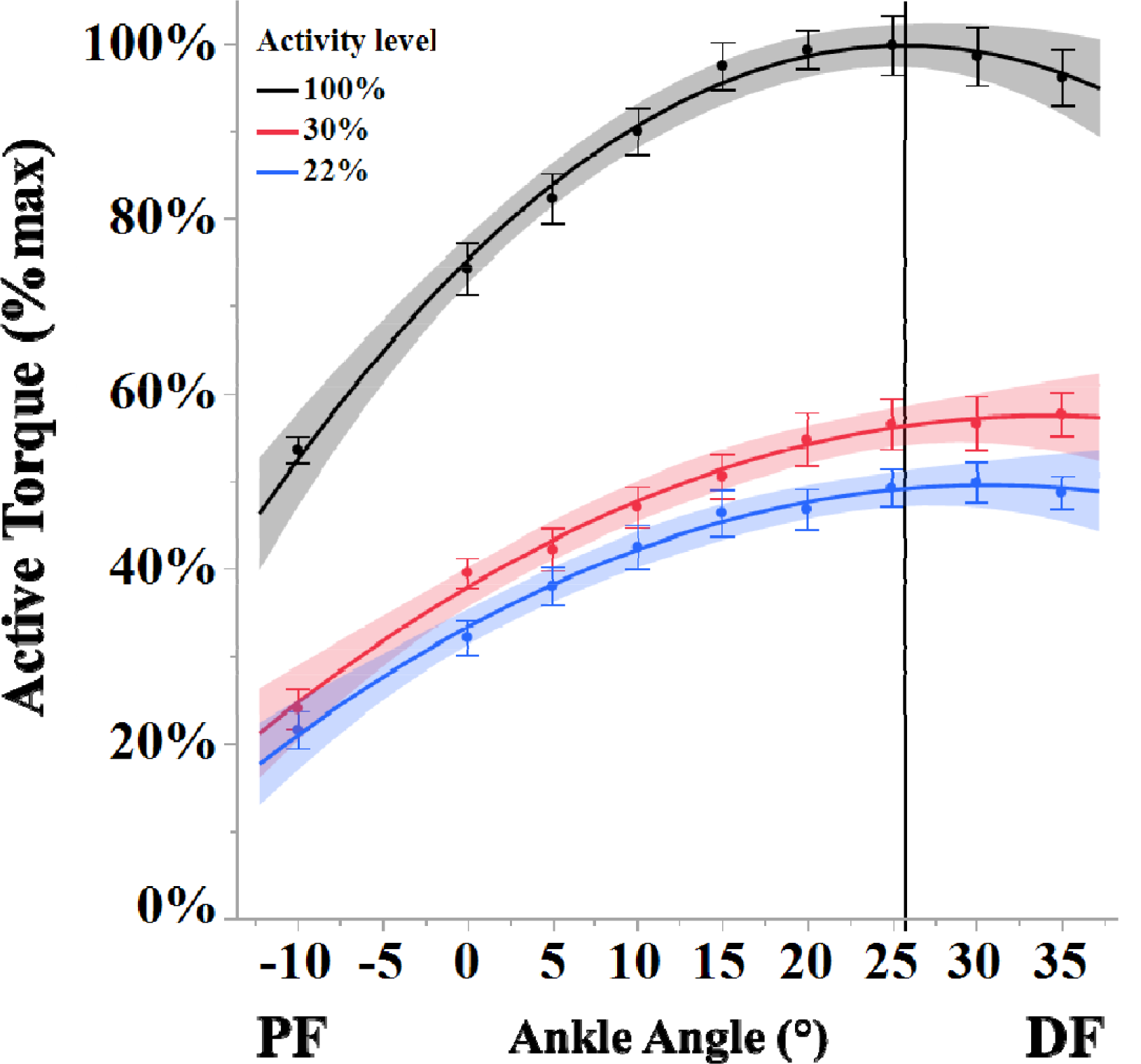
Torque-ankle angle relationships at 100%, 30%, and 22% activity levels. A binomial line is fit at each activity level and shaded regions indicate 95% confidence intervals of the fit. From the binomial equations, the optimal angle at 100% was calculated (vertical line). No optimal angle was calculated for 22% and 30% activity because no descending limb was produced from the data. From these measurements, a change in the torque-angle curve towards more dorsiflexed positions with decreasing activity level is visible. Data is normalized to the maximum torque produced during MVCs (100%). Error bars of data points are s.e.m.

**Figure 4:**
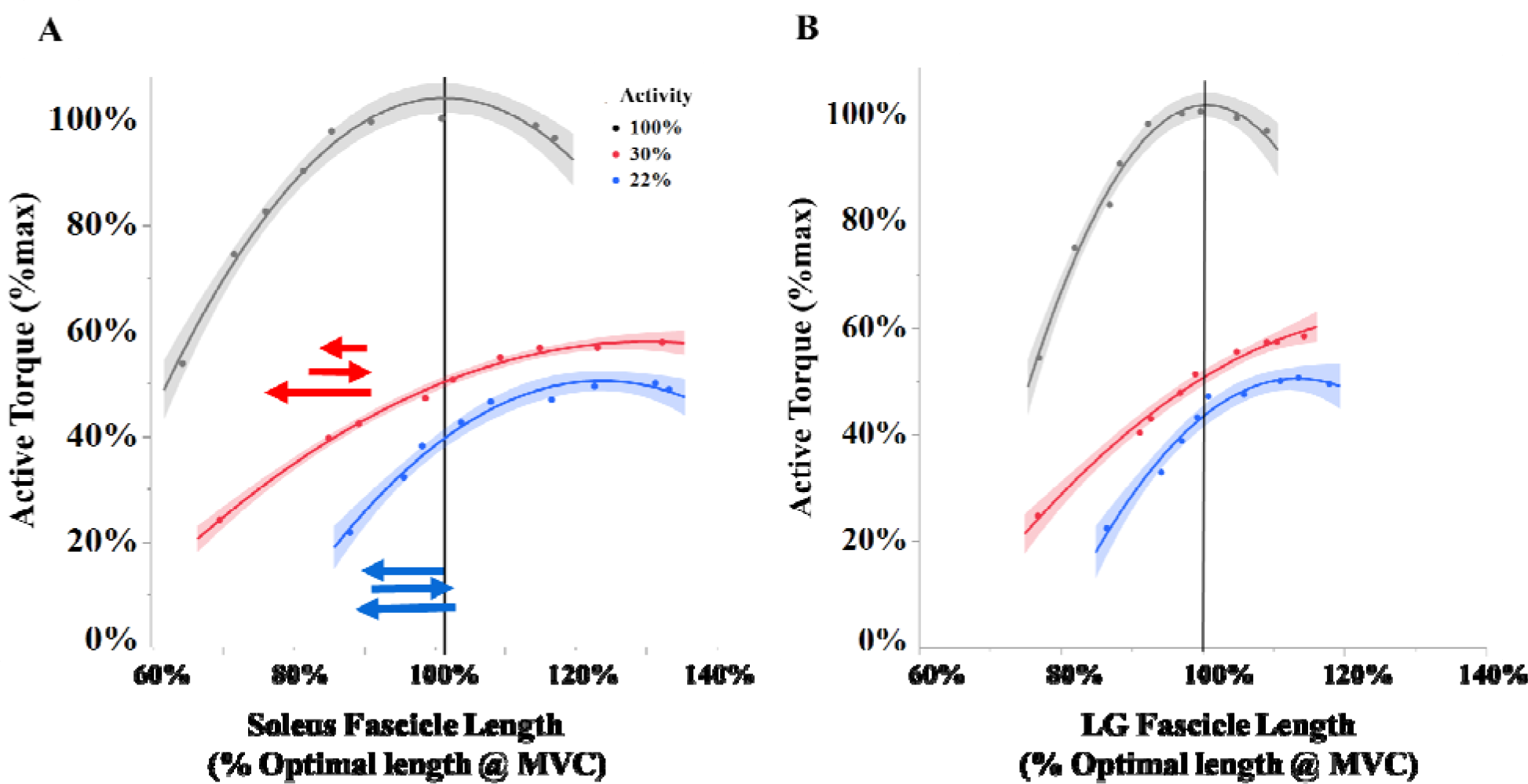
Torque-fascicle length relationships for SOL (A) and LG (B) at 100%, 30%, and 22% activity levels. A binomial line is fit at each activity level and shaded regions indicate 95% confidence intervals of the fit. From the binomial equations, the optimal fascicle length at 100% was calculated (vertical line). No optimal fascicle length was calculated for 22% and 30% activity because no descending limb was produced from the data. From these measurements, a shift of optimal fascicle length towards longer lengths with decreasing activity level is visible for both muscles. Note that because a descending limb is not present for any submaximal relationship, the exact location of the optimal fascicle length is not known. Arrows indicate previously measured fascicle operating lengths during walking during early (top arrow), mid (middle arrow), and late (bottom arrow) stance while walking without (red; (Cronin et al., 2013; Ishikawa et al., 2005; Rubenson et al., 2012)) and with (blue; (Jackson and Collins, 2015)) an ankle exoskeleton. Data is normalized to the maximum torque produced during MVCs (100%) and the corresponding fascicle length.

**Table 2.** Results of ANOVA model to describe the relationship between ankle angle, activity level, and fascicle length on active torque. τ = torque, SOL = soleus, LG = lateral gastrocnemius, interaction = angle × activity level interaction, *F* = F statistic, *P* = P-value. * indicates significance (*P* < 0.05). Note: individual random response added to the model, and significant for all tests (Wald test, P < 0.05). SOL = soleus, LG/MG = lateral and medial gastrocnemius.

|  | Angle |  | Activity Level |  | Interaction |  | Fascicle length |  |
| --- | --- | --- | --- | --- | --- | --- | --- | --- |
| | $F$ | $P$ | $P$ | $F$ | $P$ | $F$ | $P$ | $F$ |
| τ- SOL fascicle length | 38.93, <0.001* | | 420.55, <0.001* | | 1.77, 0.04 | | 15.69, <0.001* | |
| τ- LG fascicle length | 21.41, <0.001* | | 355.75, <0.001* | | 0.76, 0.07 | | 0.02, 0.89 | |

### Torque-Fascicle Length Relationship

For both SOL (Fig. 4A) and LG (Fig. 4B), decreasing their activity level increased their fascicle lengths at a specific ankle angle (Tukey HSD, P < 0.05; Fig. 4; Suppl. Fig. 5). In general, the shape of the torque-fascicle length relationship covered a smaller fascicle length range for the LG than the SOL, but the trends observed for decreasing the activity level are similar. Like the torque-ankle angle curves, while the 100% activity level relationship had an ascending, plateau and descending limb, the submaximal activity levels displayed only an ascending and plateau region, and no descending limb. Thus, for both SOL and LG, no descending limb was physiologically possible (with our experimental design, which involved an extended knee). For both muscles, the ascending limb extended to longer fascicle lengths during the 22% compared to the 30% activity level. This qualitative observation suggests a general shift of the 22% activity level curves towards longer fascicle lengths compared to the 30% condition. Based on this data, decreasing activity level changes the torque-fascicle length relationship for SOL and LG, but because only ascending and plateau regions are visible, it is difficult to estimate optimal fascicle length for the submaximal conditions.

## Discussion

The objective of this study was to evaluate how net ankle joint torque-fascicle length relationships of the major plantar flexors were affected during voluntary fixed-end contractions at two submaximal activity levels to inform us about human-machine interactions when wearing a passive assistive device during walking. The activity levels of 22% and 30% MVC replicate those observed during the mid-stance phase of level ground walking at a self-selected speed with or without an assistive lower-limb exoskeleton, respectively. Our major observation was that with decreasing voluntary muscle activity levels, the torque-ankle angle and torque-fascicle length relationships changed shape, with the curves shifting towards more dorsiflexed ankle positions and longer fascicle lengths, respectively. The shifting curves abolished the descending limb of the torque-ankle angle and torque-fascicle length relationships. The descending limb would presumably exist at ankle levels not physiologically possible > 35° with an extended knee.

### Torque-fascicle length relationship

During 100% activity, both SOL and LG fascicles achieved an ascending limb, a plateau region with an optimal fascicle length at ~25°, and (albeit short) a descending limb. In contrast, SOL and LG fascicles only produced an ascending limb and plateau region at the SOL-matched 22% and 30% activity levels. The ascending limb of the torque-fascicle length curve was shifted towards longer lengths for the 22% compared to the 30% activity levels. This shift was greater at more plantar flexed angles; for example, at an ankle angle of −10°, SOL fascicle length was shifted to ~20% longer fascicle lengths at 22% compared to 30% activity. Similar to the torque-ankle angle relationship, the torque-fascicle length relationship can be assumed to have a descending limb at ankle angles >35°, which for our participants were not physiologically possible with an extended knee.

Although we cannot precisely locate the optimal fascicle lengths, based on the graphs, we can see that the SOL and LG exhibit activation-dependence of optimal fascicle length; decreasing activity level increased SOL and LG fascicle lengths. The underlying mechanism to explain this phenomenon is a current hot topic of research (Holt and Williams, 2018; MacIntosh, 2017). Activation-dependence of optimal length, often called length-dependent activation, is an umbrella term that covers subcellular and MTU level mechanisms that are not necessarily mutually exclusive (Holt and Williams, 2018). The first mechanism is subcellular in nature and often observed in *in vitro* muscle preparations, where at a relative level of submaximal Ca^2+^, more force is produced than expected (Hessel et al., 2019; Stephenson and Williams, 1982; Yang et al., 1998). The exact molecular mechanism for this property is not yet clear, but seems to be connected to molecular changes of sarcomeric proteins (Hessel et al., 2019; Konhilas et al., 2002). Outside of subcellular mechanisms, an important mechanism responsible for length-dependent activation is associated with MTU compliance (Holt and Williams, 2018; MacIntosh, 2017) and was studied by others in the human vastus lateralis (Austin et al., 2010; de Brito Fontana and Herzog, 2016; Ichinose et al., 1997; Lloyd and Besier, 2003) and in the triceps surae (Clark and Franz, 2019). When *in vivo* MTUs are tested, there is a large activation-dependent shift in optimal triceps surae MTU length caused primarily by MTU compliance. During contraction, muscle fascicles produce force and as activation increases, the muscle fascicles shorten further, subsequently increasing the strain of the in-series Achilles tendon. Thus, the amount of Achilles tendon strain for a given joint configuration is related to the amount of muscle activity (and muscle force). The overall result is that reduced activity levels lead to a shift in the optimal ankle angle and optimal fascicle length towards more dorsiflexed ankle angles (and MTU lengths) and longer fascicle lengths, respectively.

Fascicle force should also be altered by fascicle shortening during activation because of the history-dependent property of residual force depression (Chen et al., 2019), which is a depressed force relative to the force observed following no shortening. Because the amount of muscle activity is relatively proportional to the amount of muscle shortening and muscle force (and subsequent work), which is proportional to the amount of residual force depression (Granzier and Pollack, 1989; Herzog et al., 2000), the amount of residual force depression will also be activation-dependent. Raiteri et al. (2019) recently demonstrated that during voluntary fixed-end contractions, the anterior tibialis exhibited residual force depression from activation-induced fascicle shortening, so we would also predict residual force depression to arise during the substantial fascicle shortening observed in the SOL and LG (>20% shortening from 0% to 30% activity). Although we assume that residual force depression is less for the 22% compared with the 30% activity level, we are not sure if these potential changes in residual force depression are biologically relevant in the context of human-exoskeleton interactions. Future studies similar to those of Raiteri et al. (2019) will be needed to study this phenomenon.

Are the changes to torque-fascicle length curves between 30% and 22% activity levels physiologically relevant? To illuminate this, we overlaid literature values of fascicle lengths during the stance phase of walking onto our torque-fascicle length figures for SOL (Fig. 4A, arrows; LG data was not readily available). For unassisted walking, we combined ultrasound-measured SOL fascicle data from several reports (Cronin et al., 2013; Ishikawa et al., 2005; Rubenson et al., 2012). For walking with an exoskeleton, we used model-derived estimates of fascicle lengths (Jackson and Collins, 2015). This comparison (see Fig. 4A) indicates that with decreasing muscle activity, operating fascicle lengths during walking increase. Furthermore, this increase in fascicle length seems to track closely with the shift of the torque-fascicle length curve, suggesting that they are coupled. Finally, the ascending limb during 22% MVC is steeper than that during 30% MVC, which suggests that during the crucial powered plantarflexion of stance, muscle force decreases faster during fascicle shortening with lower (22% vs. 30%) muscle activity.

An activation-dependent change in torque-fascicle length properties also means that the underlying sarcomeres are likely changed by a similar amount, which could have further consequences for force production. This data can be extrapolated to estimate the sarcomere lengths of SOL and LG at fascicle lengths and activity levels typical during stance. Previous work modeled sarcomere lengths during plantar flexor MVCs (100% activity) and reported that LG and MG sarcomere lengths are < 2.1 μm (Maganaris, 2004), much shorter than that required for maximal overlap and force production (~2.7 μm in humans (Gollapudi and Lin, 2009)). If we assumed a linear increase in sarcomere length to fascicle length, then based on our data, the operating sarcomere length becomes longer along with fascicle length at 30%, and even longer at the 22% activity level. This has the potential to move the sarcomeres to more favorable filament overlap for force production. However, it is important to consider that measuring sarcomere length *in vivo* during voluntary contractions still remains a difficult task. Technology to image and measure sarcomere lengths during *in vivo* muscle contraction is advancing rapidly (Lichtwark et al., 2018; Sanchez et al., 2015) and so direct measurements of SOL and LG sarcomere lengths during walking will hopefully be possible to confirm our estimates in the future.

Although the activity-dependent changes to torque-fascicle length properties seem relatively substantial, it is important to note that the simulated decrease in muscle activity used here was derived from data where the ankle exoskeleton stiffness was set at a level that minimized metabolic cost for the user (Collins et al., 2015). Thus, there are cost-benefit tradeoffs. The benefits of reduced muscle activity are most obviously the reduction in the metabolic cost (Collins et al., 2015). The costs of reduced activation are more complicated to quantify. Two modeling studies (Jackson et al., 2017; Sawicki and Khan, 2016) indicate that increasing the amount of exoskeleton torque (i.e. by increasing the spring stiffness in the Collins et al. (2015) design) changes the plantar flexor MTU mechanics, fascicle shortening velocities, and plantar flexor positive work outputs. From our work, we add that activation-dependent changes to the torque-fascicle length curves also produce consequences for SOL and LG force production. However, the associated costs of operating at different fascicle lengths at the activity levels we selected were outweighed by the benefits of reduced muscle activation. In the future, it would be interesting to reduce muscle activity further and quantify the torque-fascicle length cost-benefit properties.

### Implications for exoskeletons: collision forces

Of equal importance to MTU dynamics is the relationship between series ankle elasticity and minimizing opposite limb heel collision (Ruina et al., 2005; Zelik et al., 2014). Significant forward energy gained by powered plantarflexion can be lost during heel collision of the opposite limb, requiring an increase of positive work from muscles for the walker to maintain speed (increasing metabolic costs). Collision losses are minimized by a well-timed push-off of the trailing limb just before collision of the leading leg, which leads to the heel driving into the ground at an angle that is modeled to limit collisions costs considerably (Kuo et al., 2005; Ruina et al., 2005). Zelik et al. (2014) used a simple lower limb model to demonstrate that even small changes to plantar flexor series elasticity affects trail-leg push-off timing, altering the heel-ground collision angle and increasing collision-induced energy loss. Furthermore, because of natural increases in step length with walking speed (and presumably an increase in muscle activity), optimal series elastic stiffness remains relatively constant among walking speeds (Zelik et al., 2014). In our data, SOL fascicle lengths at 22% vs. 30% muscle activities were longer, indicating that the Achilles tendon was stretched less, which may result in a more compliant MTU, considering that force is low and the tendon may still operate in its toe region. Thus, it is possible that wearing a lower limb assistive device that reduces muscle activity has the potential to decrease in series elastic stiffness, and ultimately lead to an increase in collision force and the metabolic cost of transport. The mechanical properties of ankle exoskeletons can be modified to affect push-off timing (Galle et al., 2017), perhaps neutralizing the negative effects of reduced ankle elasticity. Collision forces have been vigorously evaluated in lower limb prostheses (Caputo and Collins, 2014; Herr and Grabowski, 2012; Houdijk et al., 2009) and need to be conducted in exoskeletons. We predict that the decrease in activity level associated with wearing lower limb assistive devices has the potential to increase collision-based forces and thereby increase the metabolic cost of walking.

### Limitations

Although we simulated muscle activity reduction when wearing an exoskeleton, it would be informative to have participants produce the true muscle activity reduction while wearing the exoskeleton. Making a comparable dynamometer test would require some protocol and apparatus modifications to secure the subject into the dynamometer without damaging the exoskeleton. Further, to induce the “off-loading” effect of the exoskeleton, the exoskeleton spring would need to be pre-loaded to a length comparable during the push-off phase of walking. This will be subject-specific and will most likely require some biomechanical analysis of the person wearing the exoskeleton while walking. Unlike the SOL, the LG is biarticular and so its fascicle length is affected by both ankle and knee angles. In this study, we maintained a constant knee angle, but during walking, the knee angle changes, altering LG fascicle length. Another study that systematically changes the knee angle is needed to understand the role of knee angle on LG fascicle lengths (and perhaps SOL fascicle lengths, due to intramuscular force transmission (Maas and Sandercock, 2010) at different activity levels. We did not measure the fascicle lengths of the MG because we only had access to one ultrasound transducer and we could not capture global three-dimensional behavior of the LG and SOL. Finally, we did not calculate fascicle forces because of the inherent difficulties in estimating force, which requires knowledge of (1) the subject-specific moment arms at different muscle forces and ankle angles (Maganaris et al., 1998), (2) the individual muscle force contributions to the net joint torque at different muscle activities and ankle angles, which is typically assumed to be fixed based on relative physiological cross-sectional area (Fukunaga et al., 1996) and force contributions from smaller muscles are often ignored (Zelik and Franz, 2017), and (3) accurate and representative pennation angles from each muscle to estimate muscle force from tendon force, which is difficult because muscle architecture is not homogeneous (Bolsterlee et al., 2018).

### Conclusion

The purpose of this study was to use an *in vivo* technique to explore human-exoskeleton interactions when users wear a passive ankle exoskeleton. Of interest for this experiment was if the reduction in SOL muscle activity that occurs when wearing the exoskeleton changes the torque-ankle angle and torque-fascicle length relationships. Compared to MVC, 22% (simulated exoskeleton use) and 30% (no exoskeleton use) SOL activity levels caused torque-fascicle length curves to shift to longer lengths. This shift is similar to the modeled shift in fascicle operating length predicted from the literature (Jackson et al., 2017). While these changes likely add a cost to movement efficiency, it is still outweighed by the benefit of reduced muscle activity. A further decrease in activity level should supposedly make costs outweigh the benefits, as previously suggested (Jackson et al., 2017), but this has not been empirically shown to date. Plantar flexor force is a product of a carefully tuned neuromuscular system that minimizes the cost of force production during walking. The fundamental *in vivo* research presented here is a small step towards understanding the human-machine relationship that is critical for the development of new devices with the overall goal of enhancing exoskeleton user performance and their quality of life.

## Acknowledgements

We thank T. Weingarten for help with the experimental setup.

